# Dopamine depletion affects vocal acoustics and disrupts sensorimotor adaptation in songbirds

**DOI:** 10.1101/600874

**Authors:** Varun Saravanan, Lukas A Hoffmann, Amanda L Jacob, Gordon J Berman, Samuel J Sober

## Abstract

Dopamine is hypothesized to convey important error information in reinforcement learning tasks with explicit appetitive or aversive cues. However, during motor skill learning the only available feedback signal is typically an animal’s evaluation of the sensory feedback arising from its own behavior, rather than any external reward or punishment. It has previously been shown that intact dopaminergic signaling from the ventral tegmental area – substantia nigra compacta complex (VTA/SNc) is necessary for vocal learning in response to an external aversive auditory cue in songbirds. However, the role of dopamine in learning in the absence of explicit external cues is still unclear. Here we used male Bengalese finches (*Lonchura striata* var. *domestica*) to test the hypothesis that dopamine signaling is necessary for self-evaluation driven sensorimotor learning. We combined 6-hydroxydopamine (6-OHDA) lesions of dopaminergic terminals within Area X, a songbird basal ganglia nucleus critical for vocal learning, with a headphones learning paradigm that shifted the birds’ auditory feedback and compared their learning to birds without lesions. We found that 6-OHDA lesions affected song behavior in two ways. First, over a period of days lesioned birds systemically lowered their pitch regardless of the presence or absence of auditory errors. Second, 6-OHDA lesioned birds also displayed severe deficits in sensorimotor learning as measured by their adaptive change in pitch in response to the pitch-shifted auditory error. Our results suggest roles for dopamine both in motor production and in auditory error processing during vocal learning.

**Significance Statement:** Dopamine has been hypothesized to convey a reward prediction error signal in learning tasks involving external reinforcement. However the role dopamine plays in tasks involving self-guided error correction in the absence of external reinforcement is much less clear. To address this question, we studied the role of dopamine in sensorimotor adaptation using male Bengalese finches, which spontaneously produce a complex motor behavior (song) and are capable of modulating their behavioral output in response to induced auditory errors. Our results reveal that in addition to conveying errors in motor performance, dopamine may also have a role in modulating effort and in choosing a corrective response to the auditory error.

## Introduction

Complex organisms perform sensorimotor learning to modulate behavior in response to sensory feedback. This process uses feedback from past performances arising from either explicit reward/punishment cues (e.g. food reward, electric shocks) or from self-evaluation of the performance (e.g. hearing one’s own voice during speech or song). While prior work has taken a number of approaches to taxonomizing different forms sensorimotor learning, including distinguishing model-based and model-free learning (Wolpert et al., 1995; Mohan et al., 2011; Haith and Krakauer, 2013) and habitual versus goal-directed behavior (Balleine and O’doherty, 2010; Redgrave et al., 2010), here we focus on distinguishing it broadly into two distinct components: error-based learning that relies on self-evaluation and reinforcement learning that relies on cues from the environment (Wolpert et al., 2011). Classic studies have linked dopamine to reinforcement learning as a reward prediction error signal that conveys information about explicit rewards and punishments (Schultz et al., 1997; Glimcher, 2011). However, the question of whether dopamine is also involved in error-based learning in the absence of external rewarding or aversive cues has been harder to address. Some studies have reported deficits in error-based learning in patients with Parkinson’s disease (Paquet et al., 2008; Mollaei et al., 2013), but since Parkinson’s disease is associated with cognitive and executive deficits in addition to larger motor deficits (Lees and Smith, 1983; Cooper et al., 1991; Dubois and Pillon, 1996; Jankovic, 2008), the specific role of dopamine has been difficult to isolate.

Songbirds have emerged as an effective model system in which to study the role of dopamine in sensorimotor learning. Songbirds spontaneously produce songs hundreds of times per day. Like human speech, song is learned during development (Lipkind et al., 2013) and actively maintained by auditory feedback through adulthood (Sakata and Brainard, 2006, 2008; Sober and Brainard, 2009; Kuebrich and Sober, 2015). Additionally, songbirds have a well-defined neural circuitry with separate nuclei dedicated to song production and song learning (Sohrabji et al., 1990; Scharff and Nottebohm, 1991; Brainard and Doupe, 2000). Dopaminergic neurons from the ventral tegmental area/substantia nigra pars compacta (VTA/SNc) complex innervate Area X, a basal ganglia nucleus essential for song learning, and have been hypothesized as a way for auditory error information to enter the song system (Bottjer, 1993; Soha et al., 1996; Mandelblat-Cerf et al., 2014; Peh et al., 2015) (see Fig. 1). More recently, it has been reported that birds displayed deficits in vocal reinforcement learning when dopaminergic innervation of Area X was reduced (Hoffmann et al., 2016; Hisey et al., 2018). Furthermore, neural recordings of dopaminergic neurons revealed prediction error type responses when birds were required to avoid aversive auditory perturbations while singing (Gadagkar et al., 2016) and pitch contingent optical stimulation of dopaminergic terminals in Area X evoked changes in the pitch of the birds’ song (Hisey et al., 2018; Xiao et al., 2018). However, in all of the above studies, an external reinforcement cue or equivalent optical stimulation was present, leaving open the question of whether dopamine is involved in error-based learning that relies exclusively on self-evaluation of motor behavior.

**Figure 1:**
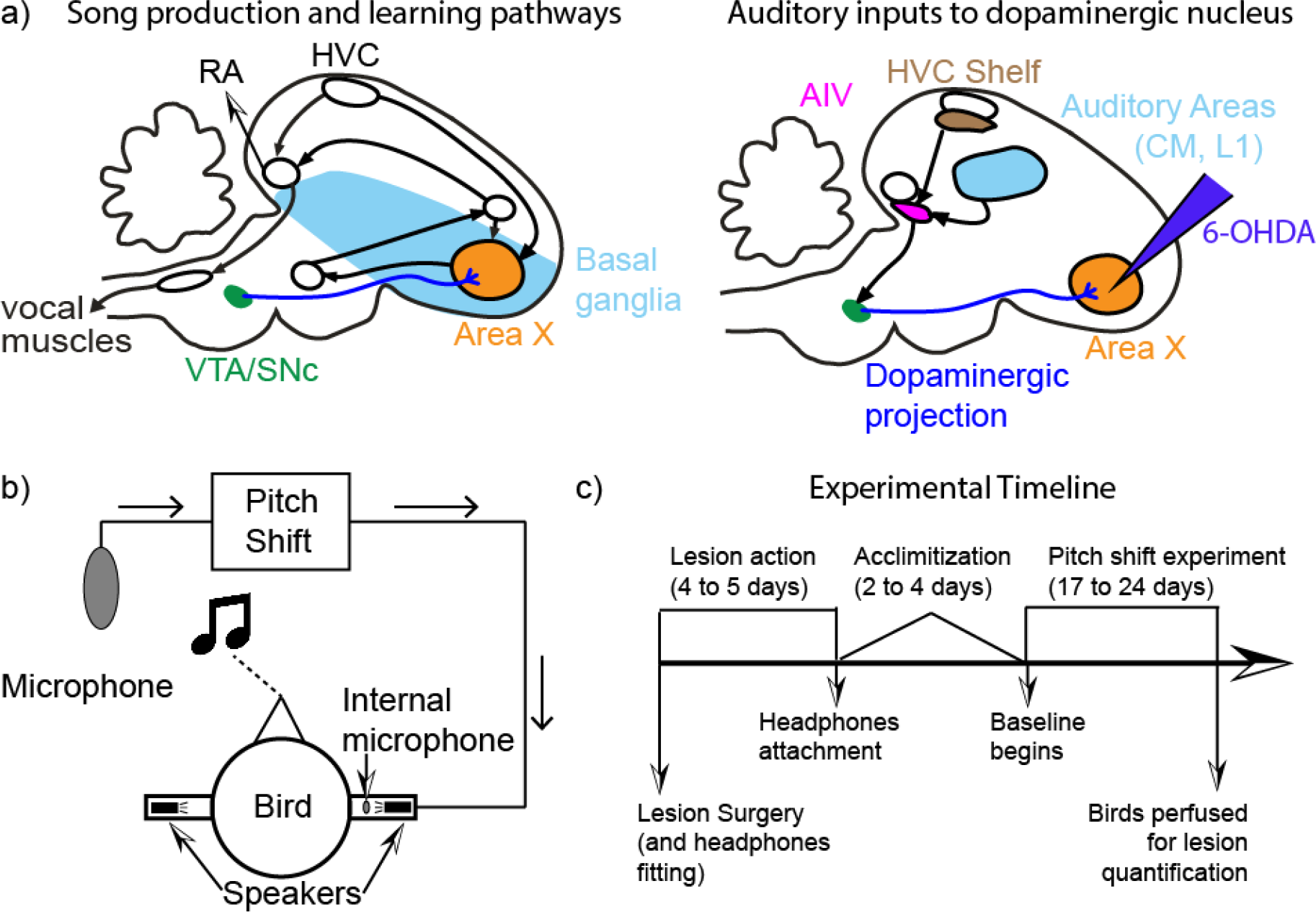
Songbird neuroanatomy and experimental design. a) A theory for the role of dopamine in sensorimotor learning in songbirds. The left panel shows the brain nuclei in the songbird primarily involved in song production and learning. Area X, a songbird basal ganglia nucleus critical for song learning, receives dense dopaminergic projections from the VTA/SNc complex. The right panel shows the nuclei involved in auditory processing in the songbird. One of the known pathways for auditory information to influence song learning is through the dopaminergic projections to Area X. We target these projections when we perform 6-hydroxydopamine (6-OHDA) lesions into Area X as depicted. b) A schematic for how the custom-built headphones introduce a pitch shifted auditory error to the birds. Briefly, a cage microphone records all sounds made within the cage and sends it through a pitch shifting program which is subsequently played back to the bird through miniature speakers attached to the headphones. The headphones also have an internal microphone to record output from the headphones speakers and to calibrate sound intensity. c) A detailed timeline for each of our experiments (see Materials and Methods).

We tested the role of dopamine in error-based learning by selectively lesioning dopaminergic terminals in Area X using 6-hydroxydopamine (6-OHDA). Since the cell bodies of dopaminergic neurons in VTA/SNc that innervate Area X are intermingled with those projecting to the rest of the songbird basal ganglia (Person et al., 2008), we injected 6-OHDA directly into Area X to avoid introducing general motor or song production deficits. We fitted the birds with custom-built headphones through which we introduced a shift in pitch (either upwards or downwards) of the bird’s auditory feedback (Sober and Brainard, 2009; Hoffmann et al., 2012) to measure how birds changed their pitch over time in response to this induced sensory error and how self-guided error correction was affected by dopamine manipulations.

## Materials and Methods

All animals used for this study were adult (range of ages: 105 to 217 days post hatch; median age: 141 days post hatch) male Bengalese finches (*Lonchura striata* var. *domestica*). Throughout the study, the animals were housed in isolated sound attenuating chambers (referred to as sound boxes) on a 14-10 hour light-dark cycle. All singing analyzed for this paper was undirected song, i.e., songs sung in the absence of a female. All procedures were approved by Emory University’s Institutional Animal Care and Use Committee.

### Experimental design and Statistical Analysis

We designed a group comparison study to test the role of dopamine in sensorimotor adaptation. We performed pitch shift experiments on 6 unlesioned birds (3 each for upward shifts and downward shifts) and 8 lesioned birds (4 for upward pitch shift and 4 for downward pitch shift). As detailed below, virtual auditory feedback through the headphones was delivered almost in real time and was meant to replace the natural auditory feedback that birds would otherwise receive. All pitch shifts were 1 semitone in magnitude (equally split between +1 and −1 semitone shifts). Each experiment consisted of 3 days of baseline (unshifted auditory feedback through headphones) followed by 14 days of pitch shifted auditory feedback. At the end of the shift period, we turned off the shift in pitch (i.e. set the pitch shift to zero semitones as in the baseline epoch) and recorded the birds’ activity for 6 to 7 days. During this period, unlesioned birds typically reverse the effects of the pitch shift (Sober and Brainard, 2009). We refer to this period as “washout.” Washout data were collected for all 6 unlesioned birds. Due to technical difficulties associated with keeping the headphones attached for extended periods of time, washout data was collected for only 4 out of the 8 lesioned birds (2 for upward pitch shifts and 2 for downward). In addition, we performed control experiments with 2 unlesioned birds fitted with headphones and no pitch shift and 8 lesioned birds without any pitch shifts (5 with headphones and zero pitch shift throughout; 3 with no headphones). To minimize the number of animals we used, our unlesioned bird group consisted of data reanalyzed from Sober and Brainard, 2009. Furthermore, since we showed previously (Hoffmann et al., 2016) that animals injected with saline instead of 6-OHDA were statistically indistinguishable from unlesioned birds, we did not include a saline injected control group in this study. Note that of the 8 birds whose data were reanalyzed from Sober and Brainard (2009), the raw data for 2 animals – the unlesioned birds with no headphones shift – were unavailable. However we were able to extract the daily mean pitch values from each animal’s data from an eps version of the original figure summarizing the data. The resulting figure that shows the mean change in pitch and error bars for the group was produced from the 2 data points for each day.

For our lesioned group, we reduced the dopaminergic innervation of Area X (Fig. 1), a song specific nucleus of the basal ganglia, using 6-hydroxydopamine (6-OHDA) microinjections as described in detail previously (Hoffmann et al., 2016). Briefly, we used stereotactic surgeries to target Area X with a 4 × 3 grid of microinjections of 6-OHDA (see 6-OHDA lesions below). Following 6-OHDA surgery, the birds were allowed to recover in their sound boxes for 4 to 5 days which also served as a period to allow the 6-OHDA to cause degeneration of striatal innervation (Jeon et al., 1995). Subsequently, the headphones (Hoffmann et al., 2012) were fitted to the birds and set to initially provide unshifted auditory feedback (zero pitch shift). Following headphones attachment, the birds typically did not sing for 2 to 4 days (see Fig.1c for a timeline schematic). Once they started singing again (defined as at least 30 song bouts produced over the entire day), we began recording a 3 day baseline period. Following the 3 days of baseline, the birds were recorded for 14 days during a period of shift. As described previously (Sober and Brainard, 2009; Kelly and Sober, 2014), the pitch shift was a 1 semitone shift (either upwards or downwards) played back to the bird through the headphones. The auditory feedback through the headphones was almost real-time (delay of around 10 ms) and was intended to replace the bird’s natural auditory feedback. In order to do so, the volume is set to be at least 2 log units greater in sound intensity than the bird’s own feedback. For the birds that had no pitch shift through the headphones, they continued with zero shift as they were in baseline for the equivalent 14 days. Following this 14 day period, we recorded the birds’ activity for 6 to 7 days of washout. Owing to the difficulties of keeping the headphones attached and functional for long periods of time, we were not able to collect washout data for every animal. Analysis of washout data was therefore necessarily limited to data from birds that did have data collected for the washout period.

Note that one of our 6-OHDA lesioned birds in the −1 semitone shift group was subjected to an extended baseline period of 6 days rather than the 3-day period used for all other animals. Excluding data from this bird did not change any of our results significantly. Therefore all results reported include this bird, treating the last three days of baseline equivalent to days 1 through 3 of baseline for every other bird.

Birds with lesions that were not fitted with headphones were returned to their sound boxes post-surgery and were recorded for the duration of the experiment. In this case, since they did not have a break in singing due to placement of fully assembled headphones, the baseline was defined as days 6 through 8 post lesion and the “shift” period was defined as day 9 through 22 post lesion to keep the timelines comparable between groups.

The statistical analyses used in this manuscript are detailed in their corresponding subsections in the Methods. We used a two-sample KS test to quantify dopamine depletion effects of the 6-OHDA microinjections (see Image and Lesion Analysis below). For our pitch quantification, we reported direct probabilities using bootstrapping and these are detailed in the Error quantification and Hypothesis testing with Bootstrap sections below. Since bootstrapping reports Bayesian probabilities, we verified our results with frequentist statistics in the form of Linear Mixed Models as is detailed in the section titled “Validating our Results with Linear Mixed Models”. We also examined correlations between lesion extent for each bird and magnitude of change in pitch using the Pearson’s correlation coefficient.

### 6-OHDA Lesions

We performed the lesions using stereotactic surgeries as described in detail previously (Hoffmann et al., 2016). Briefly, birds were anesthetized using ketamine and midazolam and positioned at a beak angle of 20 degrees below horizontal. Isoflurane was used to sustain anesthesia following the first hour of surgery. All stereotactic coordinates were relative to the landmark Y_0_, the posterior border to the divergence of the central sinus in songbirds. Small craniotomies were performed above the coordinates AP 4.75 to 6.4; ML 0.75 to 2.3 on both sides (all coordinates are in mm). 6-OHDA (Tocris; conjugated with HBr) was injected bilaterally in a 4 × 3 grid at AP coordinates 5.1, 5.5, 5.9 and 6.3 and ML coordinates 0.9, 1.55 and 2.2 with a DV coordinate between 3.08 and 3.18 from the surface of the brain. Additionally, for some birds, there was one final injection at AP 4.8, ML 0.8 and DV 2.6 from the surface of the brain. Each injection injected 13.8 nL of 6-OHDA in the slow setting using a Drummond Scientific (Broomall, PA) Nanoject II auto-nanoliter injector.

### Headphones attachment and assembly

The methodology is described in detail in (Hoffmann et al., 2012). Briefly, each set of headphones was custom-fit to an individual bird under anesthesia. If attached on a bird that also had a 6-OHDA lesion, both lesion and headphones fit adjustment were performed back-to-back in the same surgery. Once the headphones had been successfully fitted for the bird, the electronics (a speaker on each side and a miniature microphone on one side to record headphones output and calibrate volume) were assembled offline. The fully assembled headphones were then refitted to the bird 4-5 days post-surgery. We used a flexible tether with a commutator to power the headphones and read the electronic signals.

### Histology

Following the end of the experiment, headphones were removed and the birds were deeply anesthetized with ketamine and midazolam before performing perfusions using 10% formalin. The brains were postfixed overnight in formalin and then cryoprotected in 30% sucrose for 1 to 4 days prior to slicing into 40 µm sections on a freezing sliding microtome. Alternating sections were either immunoreacted with tyrosine hydroxylase antibody and visualized with diaminobenzidine (TH-DAB) or Nissl stained. TH-DAB was used to quantify the extent of lesions in the 6-OHDA birds, while Nissl was used to verify that there had been no necrosis and to assist in identifying boundaries of Area X in adjacent TH-DAB sections. For the TH-DAB reaction, all incubations were carried out on a shaker at room temperature and all chemicals were dissolved in 0.1M phosphate buffer (PB) unless otherwise noted. Fixed sections were treated sequentially with 0.3% hydrogen peroxide to suppress endogenous peroxidases and 1% sodium borohydride to reduce exposed aldehydes and improve background staining before incubating overnight in a tyrosine hydroxylase antibody solution (Millipore Cat# MAB318, RRID:AB_2201528, 1:4000; 0.3% Triton X-100; and 5% normal horse serum). Tissue was then incubated in biotinylated anti-mouse secondary antibody (Vector Laboratories Cat# BA-2000, RRID:AB_2313581, 1:200 and 0.3% Triton X-100) followed by avidin-biotin-complex (ABC) solution (Vector Laboratories Cat# PK-4000, RRID:AB_2336818). Tissue was exposed to DAB solution (Amresco E733; 5 mg DAB per tablet; 2 tablets in 20 ml of purified water) for approximately 5 min. Sections were mounted, air-dried, delipidized with ethanol and citrisolv, and coverslipped with Permount (Fisher scientific, SP15-500). For the Nissl stained sections, Nissl stain was applied on mounted, air-dried tissue, which was delipidized with ethanol and citrisolv, and coverslipped with Permount. Stained sections were imaged using a slide scanner (Meyer Instruments PathScan Enabler IV; 24 bit color, 7200 dpi, “sharpen more” filter, brightness, and contrast level 50) and the resulting images were analyzed using ImageJ (RRID:SCR_003070).

### Image and Lesion Analysis

TH-DAB stained sections were used for lesion quantification by analysis through a custom written macro in ImageJ. The analysis was based on a metric of optical density described in detail in (Hoffmann et al., 2016). Briefly, the macro allowed us to demarcate the boundary of Area X in every section that it is present. We also used a 0.5 mm circle to mark a section of representative striatum outside of Area X in the same section. We then defined the quantity optical density ratio (OD ratio) as the ratio between the optical density of Area X in the section to that of striatum in the section as follows:

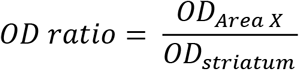

Since Area X is typically stained darker by TH-DAB than the surrounding striatum (Bottjer, 1993; Soha et al., 1996; Hoffmann et al., 2016), we used the cumulative distribution of the optical density ratio saline injected birds to define our threshold for lesions. Any section in our group of 6-OHDA lesioned birds with an OD ratio less than the 5^th^ percentile of the saline injected birds sections counted towards the overall proportion of lesioned sections. Additionally, we used a two-sample Kolmogorov-Smirnov test to test whether the lesioned and saline populations were indeed drawn from separate distributions. We also used the threshold procedure described above to quantify lesion extent for individual animals. We then asked whether lesion extent was significantly correlated with vocal behavior metrics such as baseline variance, change in variance from baseline to end of shift and change in pitch at the end of shift.

### Pitch Quantification

All our analysis was performed using an extracted value of pitch for every instance in which a bird sings a particular syllable. Briefly, birds have multiple syllables within their song and they typically repeat their song hundreds of times per day during the course of the experiment. We call each time they sing a particular syllable an iteration of that syllable. We restricted our analysis to roughly 30 song files per day between 10 am to 12 pm and have shown earlier that the choice of time window does not qualitatively affect our results (Sober and Brainard, 2009; Hoffmann and Sober, 2014; Kelly and Sober, 2014). To quantify pitch, for each syllable we specify a time during the syllable (relative to syllable onset) during which the syllable is relatively flat and clear in the frequency vs time space and can be reliably quantified across iterations across days. The pitch we extract represents a weighted average of the frequencies with the highest power in the lowest harmonic of the syllable. In order to make comparisons between different syllables whose base frequency can vary widely, we convert the pitches into semitones as shown below:

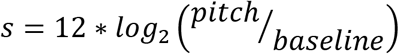

where *s* is the change in pitch in semitones, *pitch* is the observed pitch and *baseline* is the average pitch across the 3 days of baseline for that particular syllable. For all group analysis, the means reported are the means over all birds and over all syllables weighted by the proportion of times they sang each syllable. This was chosen to account for the fact that syllables that are sung more often are exposed a greater number of times to the shifted auditory feedback. Pitch quantification was performed using custom-written scripts in MATLAB (RRID:SCR_001622).

### Error quantification

For each of our groups, we had between 4 to 8 birds, each bird performed between 4 to 12 different syllables whose pitch could be quantified, and each syllable was repeated between 40 to 600 times per day. As a result, while we have several thousands of data points towards establishing the position of the mean pitch change per group for each day, the structure of the data is hierarchical and error accumulates at different levels (birds, syllables and iterations). Grouping all the data together and estimating the standard error of the mean underestimates the error by ignoring the non-independence between data points due to the hierarchical structure. On the other extreme, aggregating points and simply using individual birds or syllables does not allow us to use all of our data effectively. This is a complex problem that different studies, including our own prior efforts have used varying methods to address (Galbraith et al., 2010; Sober and Brainard, 2012; Aarts et al., 2014; Tian and Brainard, 2017). To more accurately quantify the error in our groups and better account for the variance arising from finite data samples, we use a hierarchical bootstrapping approach (Crowley, 1992; Efron and Tibshirani, 1994). In its simplest form, bootstrapping involves generating N (N = 10^4^ throughout this paper) random subsamples of the dataset by sampling with replacement from the original data and computing a metric of interest for each subsample. This results in having a distribution of the metric of interest, the 67% confidence interval of which provides an accurate estimate of the uncertainty in measurement of that metric in the original dataset (Efron, 1981, 1992; Efron and Tibshirani, 1994). For example, if one wanted to obtain the uncertainty in measuring the kurtosis of the data, one would generate bootstrap subsamples and calculate the kurtosis for each subsample. The standard deviation of the population of kurtosis values so obtained gives an accurate estimate of the uncertainty of the kurtosis in the original data. In the special case of estimating a population of means (which is the metric of interest in all instances in this paper), the uncertainty in measurement referred to above corresponds to the standard error of the mean of the dataset. However, bootstrapping by itself does not solve the problem of non-independence in hierarchical data. Crucially, to address this issue the resampling described above has to be done separately over each level of the hierarchy. This means that to generate a single subsample, we first resampled among the birds, then for each selected bird, we resampled among its syllables and finally for each syllable, we resampled among its iterations. Finally, we acknowledged that Bengalese finches can vary greatly in their syllable repertoires from one bird to the next. While all birds typically have an order of 10 syllables, some birds repeat one or two syllables with a much higher frequency than any other syllable while others represent each syllable equally. Since the bootstrapping procedure was used to calculate uncertainty of measurement due to sampling from a limited number of birds, we posited that each syllable would be equally likely in hypothetical new birds. Therefore, we set the number of iterations of a particular syllable that could occur in a bootstrapped subsample to be independent of the frequency of occurrence of that syllable in the actual data. All the data for the subsample were then combined and their mean was calculated for the subsample. Note that this procedure only applies to our estimate of measurement uncertainty (not the mean pitch values), since the means reported in the results are calculated from the actual data collected. This process was then repeated N times. In order to also account for the error in estimation of the mean of each syllable during baseline, the resampling was performed on pitch measurements recorded in hertz (Hz) and the measurements were converted to semitones just prior to calculating the mean pitch for each subsample. A similar procedure was followed for quantifying error during washout. To account for the error in estimation of pitch on the last day of pitch shift, the subtraction of the mean pitch on the final day of shift through the washout period was performed following the resampling. Our error quantification was performed using custom written scripts in MATLAB (all analysis scripts will be made available on Github post-publication).

### Hypothesis testing with Bootstrap

In addition to using bootstrapping to compute error estimates as described above, we also used a bootstrapping approach to test whether vocal pitches were significantly different across time or experimental conditions by computing direct posterior probabilities for individual hypotheses. Hence, we report our results in terms of direct probabilities of a sample being greater than or equal to another sample or fixed value in lieu of p-values. Specifically, we resample the distribution for each group and calculate the mean 10^4^ times to produce a distribution of resampled means to calculate the variance associated with having a finite number of samples.

These resampled distributions were used to compute whether the two distributions of vocal pitches were significantly different. For all instances in this paper, we use two-way tests with α = 0.05. This means that a probability is significant if the probability supporting the hypothesis, p < α/2 or if p > (1 – α/2), i.e., if p < 0.025 or if p > 0.975. In the case of computing the probability of the mean of a group being different from a constant, one can calculate the proportion of the population of bootstrapped means (as defined in Error quantification above) being greater than or equal to said constant. For example to compute the probability that the mean pitch of a particular group is significantly different from zero, one would compute the proportion of the population of bootstrapped means that are greater than or equal to zero. If this proportion is less than 0.025 then the pitch of the group of interest is significantly below zero while if the proportion is greater than 0.975 then the pitch of the group is significantly above zero.

We used a similar approach to compute significant differences between two groups of interest. In this case, we compute a population of bootstrapped means for each group. From these two bootstrapped populations, we compute a joint probability distribution between the bootstrapped means of the two groups. The null hypothesis representing no difference between the two groups would correspond to a circle centered about the unity line. Therefore, to test the difference between the two groups, we compute the volume of the joint probability distribution on one side of the unity line (including the unity line itself) to quantify the probability of one group being greater than or equal to the other group. If the probability computed is greater than 0.975, then the first group is statistically greater than the second group. Alternatively if the probability computed is less than 0.025, then the first group is statistically less than the second group. We computed multiple comparisons between groups by computing differences between 2 groups at a time and applied a Bonferroni correction to the threshold for significance. Our statistical tests were performed using custom scripts written in MATLAB which will also be made available on Github post-publication.

### Validating our Results with Linear Mixed Models

To ensure that our results were robust to our choice of error quantification and design, we also separately reported frequentist statistical tests on our results. Since our data are hierarchical (see Error quantification above), the recommended way to perform frequentist statistics on our data is through linear mixed models (Aarts et al., 2014; Aarts et al., 2015). Specifically, we built linear mixed models by using bird identity and syllable identity within a bird as variable effects and tested for significance of fixed effect factors. Concretely, our linear mixed models were of the form:

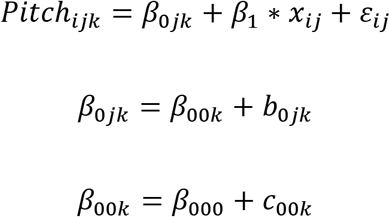

where x_ij_ refers to the condition of the shift (± 1 semitone or 0 semitone) and is the fixed effect while b_0jk_ accounts for the bird identity and c_00k_ accounts for syllable identities within a bird which are both variable effects. The code for hypothesis testing using LMMs was also done in MATLAB and will be available on Github post-publication.

## Results

We performed pitch shift experiments on 6 unlesioned birds (3 each for upward shifts and downward shifts) and 8 lesioned birds (4 for upward pitch shift and 4 for downward pitch shift). Following the end of the pitch shift, we also collected data during the “washout” period, i.e., when the pitch shift is set back to zero and the bird typically reverts its pitch back to baseline. All 6 unlesioned birds had washout data collected for 6 days following the end of shift. Of the 8 6-OHDA lesioned birds, 4 had data for washout for 7 days each (we were unable to record washout data for the other 4 lesioned animals due to technical problems associated with long-term use of the headphones). In addition, we performed control experiments with 2 unlesioned birds fitted with headphones who heard unshifted (zero pitch shift) auditory feedback and 8 birds who received 6-OHDA lesions but did not undergo any pitch shifts (see Materials and Methods for complete details).

### 1. 6-OHDA lesions reduce dopaminergic innervation of Area X

We quantified the lesion extent using a metric developed as part of our prior work (Hoffmann et al, 2016). Specifically, we used sections of Area X stained with an immunohistochemical marker (diaminobenzidine or DAB) for tyrosine hydroxylase (TH), an enzyme involved in the synthesis of dopamine and a reliable marker for dopaminergic and noradrenergic innervation (Figure 2). TH-DAB does not follow the Beer-Lambert law and varies in stain intensity even within the same animal (Van Eycke et al., 2017). As a result, quantification is typically performed between hemispheres within one section comparing a lesioned to an unlesioned hemisphere. However, we had to perform bilateral lesions for our experiments since song learning is not known to be lateralized in Bengalese finches. To quantify lesion extent, we used the fact that Area X has denser dopaminergic innervation and thus stains darker by TH-DAB than the surrounding striatum (Bottjer, 1993; Soha et al., 1996). Specifically, we quantified an optical density ratio (OD ratio) for a batch of birds that had been injected with saline into Area X (N = 4 birds; data reanalyzed from Hoffmann et al, 2016) and produced a cumulative distribution plot of the ratio across all sections for these birds. We then defined the 5^th^ percentile of that distribution as the threshold for defining lesioned sections (see Materials and Methods). When we produced a similar cumulative distribution plot of the OD ratio for all 16 of our 6-OHDA lesioned birds, around 37.5% of all sections were below the threshold defined above (Fig. 2b). This was somewhat smaller than the lesion extent for the cohort of birds in (Hoffmann et al., 2016) in which 50% of lesioned sections were below the threshold. However, the lesions were qualitatively similar between the two groups. In addition, the population of OD ratios for the 6-OHDA lesioned birds was consistently below that for the saline injected birds as verified by a two-sample Kolmogorov-Smirnov test (K = 0.3467; p = 5.75*10^−9^). We have also previously shown that such 6-OHDA lesions have no discernible effect on the existing low levels of noradrenergic innervation of Area X (Hoffmann et al., 2016).

**Figure 2:**
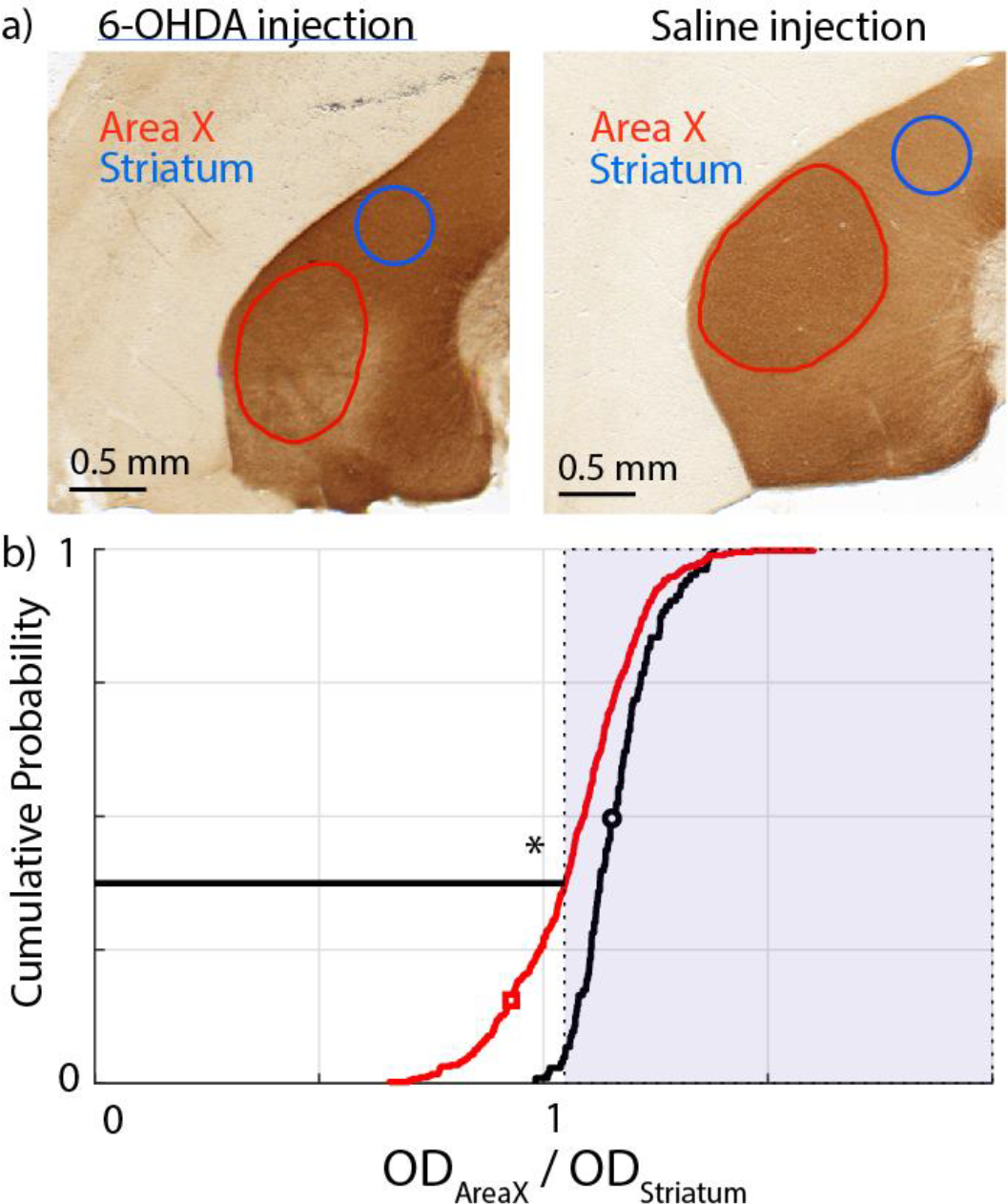
Metric for quantifying the extent of our lesions in our population of birds. We used an optical density ratio (OD ratio) between Area X and the surrounding basal ganglia (see Materials and Methods) and compared the cumulative ratios between a saline injected population (N = 4 birds) and our 6-OHDA lesioned population (N = 16 birds). a) Examples of 6-OHDA lesioned (left) and saline injected (right) sections. The red trace demarcates the Area X boundary. The blue circle is chosen to represent a uniformly stained section of the rest of the striatum. The ratio for each section is calculated as the OD ratio between these two regions. b) Cumulative distribution plots for the saline injected birds (black trace) and the 6-OHDA lesioned birds (red trace). The shaded portion represents ratios that are greater than the 5^th^ percentile for the saline injected birds. By this metric, 37.5% of all 6-OHDA lesioned sections have a smaller OD ratio. The black and red symbols correspond to the examples shown in a). The * represents a statistically significant difference between the red trace and the black trace (KS test; p<0.05; see Results for full description).

### 2. 6-OHDA lesioned birds reduce pitch even in the absence of auditory error

We showed earlier that in unlesioned animals, the headphones do not cause changes in vocal pitch in the absence of any shifts in feedback pitch (Sober and Brainard, 2009). As shown in Figure 3a, the mean pitch across days 12 through 14 of the experiment for these birds was found to be 0.02 ± 0.07 semitones (all measures of mean pitch reported are mean ± SEM). Since this particular dataset only consists of the 6 data points shown in Figure 3a, it did not make sense to perform a bootstrap analysis (here SEM is measured across 6 data points; see Materials and Methods). Instead we used a one sample t-test and found that this distribution was not significantly different from zero (t = 0.35; df = 5; p = 0.74).

**Figure 3:**
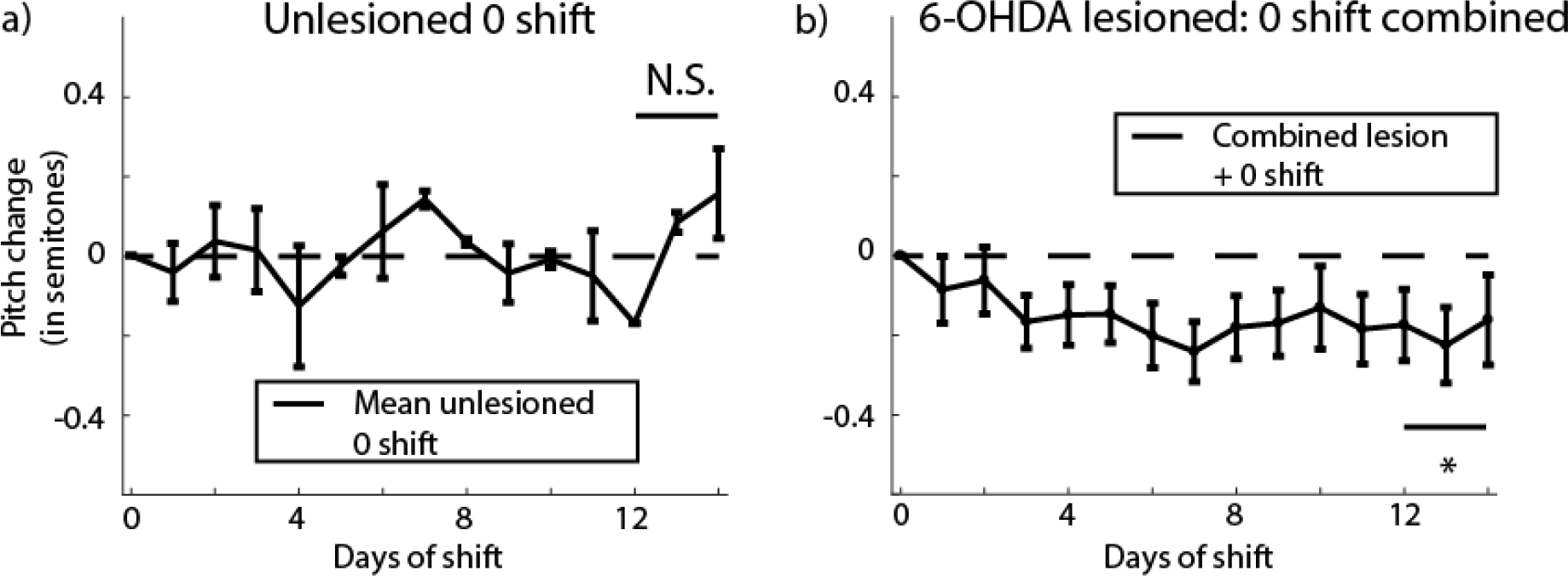
Quantifying the effect of headphones without any pitch shifts on the average change in pitch of the bird with or without lesions. a) Mean change in pitch of song for 2 unlesioned birds with headphones but no shifts through the headphones (analyzed from data extracted from Supp. Fig. 6 from Sober and Brainard, 2009). b) Mean change in pitch for 6-OHDA lesioned birds combining both birds with headphones but no shift in pitch (N = 5 birds) or without headphones (N = 3 birds) for a total of 8 birds. The group averages for the two groups and the individual traces for all 8 birds is shown in Figure 3-1. N.S. represents “not significantly different from zero” while the * represents a significant difference when comparing the last 3 days of shift combined from zero (p<0.05).

**Figure 3-1:**
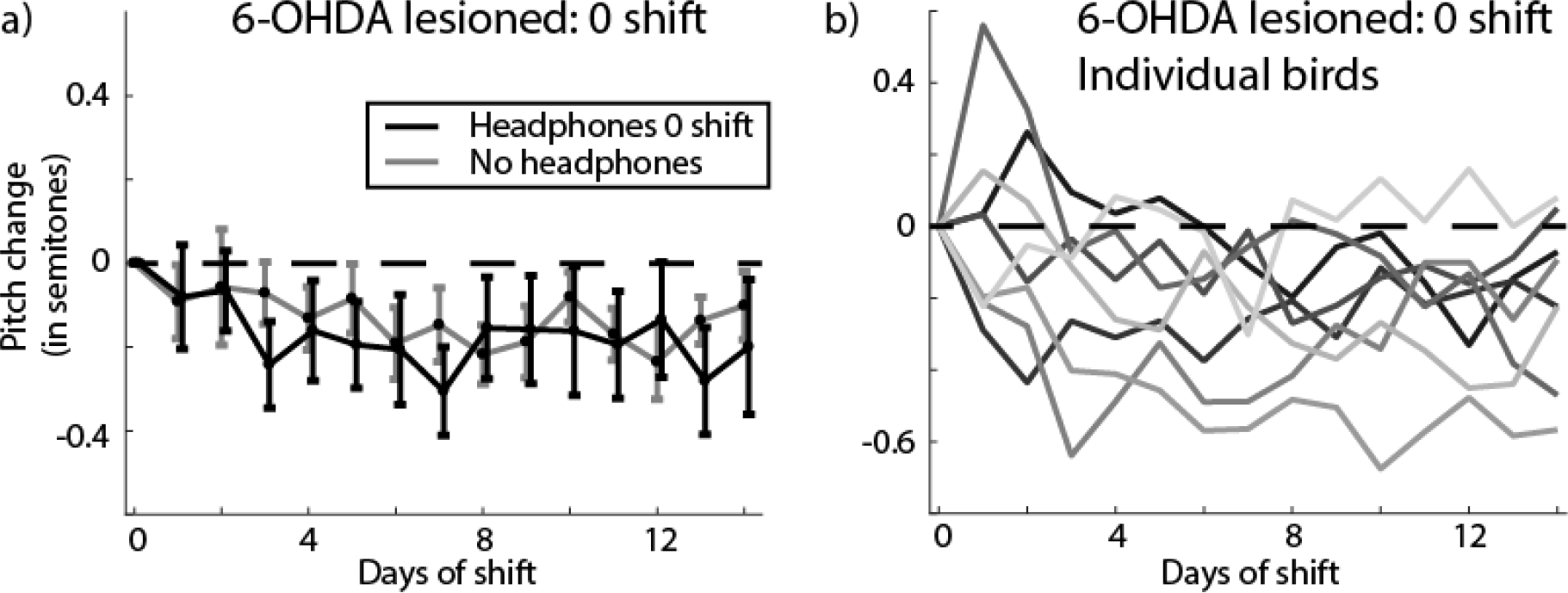
a) Mean change in pitch for 6-OHDA lesioned birds either with headphones but no shift in pitch (black trace; N = 5 birds) or without headphones (gray trace; N = 3 birds). b) Mean change in pitch for individual lesioned birds subjected to zero pitch shift either with or without headphones.

Data from 8 birds with 6-OHDA lesions but without any pitch shift revealed an unexpected systematic lowering of vocal pitch after dopamine depletion. Of those, 5 birds had headphones that conveyed unshifted auditory feedback (i.e. no pitch shift) and 3 birds had no headphones attached. When we analyzed the mean pitch change for each day for these two groups, we found them within error bars of each other for all 14 days of the experiment, and their pitch change across days 12 through 14 (−0.20 ± 0.14 with headphones; −0.16 ± 0.06 without headphones) were statistically indistinguishable (probability of resampled mean pitch with headphones greater than that without headphones was p = 0.098; see *Hypothesis testing with Bootstrap* in *Materials and Methods*). As a result, we combined the data from the 2 groups for the remainder of our analyses (the means for individual groups and traces for individual birds are shown in Fig. 3-1). The resulting mean shift in pitch during the course of the experiment is shown in Figure 3b. The overall shift in pitch over days 12 through 14 for this combined group was −0.19 ± 0.08 semitones. This decrease in pitch was statistically significant (probability of resampled mean pitch greater than or equal to zero was p = 0.0029), demonstrating, unexpectedly, that 6-OHDA lesions of Area X impacted song production by reducing the average pitch over time even in the absence of pitch-shifted auditory feedback.

### 3. 6-OHDA lesioned birds do not respond adaptively to pitch-shifted auditory error

In unlesioned animals, birds respond to a pitch shift through the headphones in an adaptive manner. Specifically, when subjected to a +1 semitone pitch shift through the headphones, the unlesioned birds compensate adaptively by lowering their pitch (mean pitch change over days 12 to 14 for N = 3 birds was −0.40 ± 0.07 semitones; blue trace, Fig. 4a; probability of resampled mean pitch greater than or equal to zero was p < 10^−4^; limit due to resampling 10^4^ times) and when subjected to a −1 semitone shift in pitch, the unlesioned birds increase their pitch (mean pitch change over days 12 to 14 for N = 3 birds was 0.36 ± 0.11 semitones; red trace, Fig. 4a; probability of resampled mean pitch greater than or equal to zero was p = 0.9996, recall that in our bootstrapping analysis we conclude that distributions are significantly different if the probability that one is greater than or equal to the other is less than 0.025 or greater than 0.975; see Methods; traces for individual birds are shown in Fig. 4-1a). The result of plotting adaptive change in pitch (inverting y-axis for +1 semitone shift birds) for unlesioned birds is shown in Figure 4c (black trace). A direct comparison between the populations of −1 semitone shift and +1 semitone shift birds revealed a complete non-overlap among posterior distributions of sampled means (probability of resampled mean pitch for +1 semitone shift greater than or equal to that for −1 semitone shift was p < 10^−4^; limit due to resampling 10^4^ times). This resampling-based analysis reaffirms our initial finding (Sober and Brainard, 2009) that unlesioned birds respond adaptively to pitch-shifted auditory errors and compensate accordingly for them, despite the fact that this earlier paper did not take into account the hierarchical nature of the data and the resulting propagation of uncertainty when computing statistical significance.

**Figure 4:**
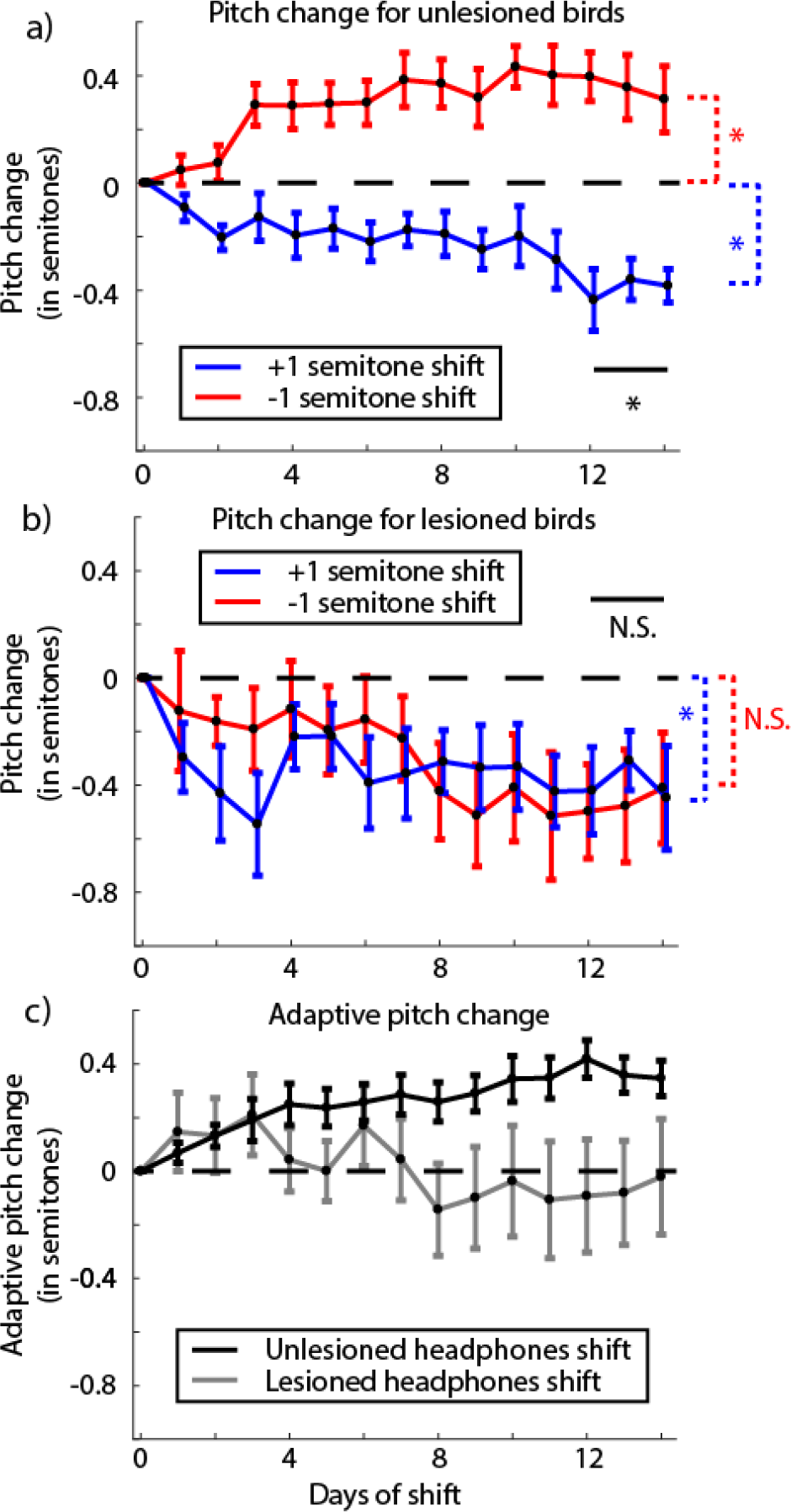
Change in pitch in response to pitch shift errors through the headphones in unlesioned and 6-OHDA lesioned birds. a) Change in pitch from baseline over the period of pitch shift for unlesioned birds broken up by the direction of introduced shift in pitch (data reanalyzed from Sober and Brainard, 2009). The graph shows that birds increase their pitch over time in response to a downward pitch shift (red trace; N = 3 birds) and decrease their pitch to an upwards pitch shift (blue trace; N = 3 birds). Traces for individual birds are shown in Figure 4-1a. b) Same graph as in a) quantified for 6-OHDA lesioned birds (N = 4 birds for each trace). Individual birds are shown in Figure 4-1b. c) Adaptive change in pitch (see Results) for unlesioned birds (black trace; N = 6 birds) and 6-OHDA lesioned birds (gray trace; N = 8 birds). For a) and b), the * and N.S. in black represent significant and not significant differences respectively between the two shift conditions while the color coded differences check difference of each group from zero (see *Results* and Table 1).

**Figure 4-1:**
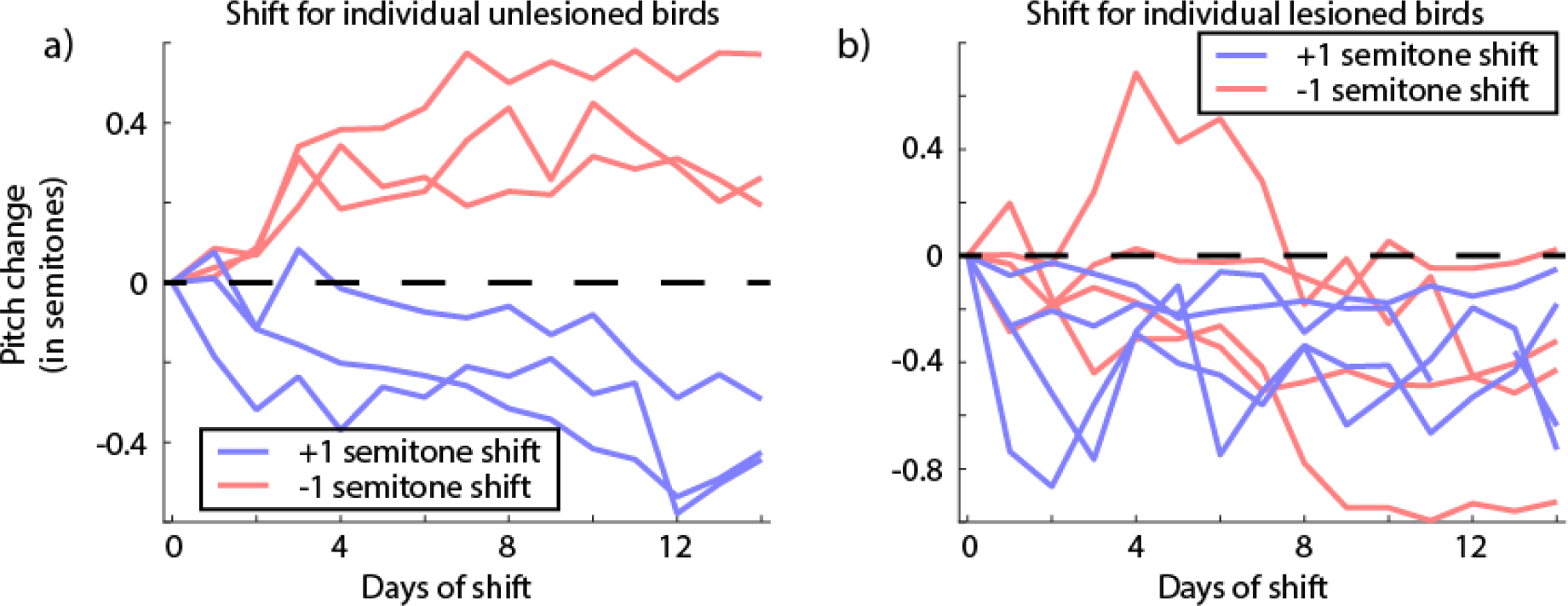
a) Mean change in pitch for individual unlesioned birds subjected to a ± 1 semitone pitch shift. b) Mean change in pitch for individual lesioned birds subjected to a ± 1 semitone pitch shift. Note that one bird subjected to a +1 semitone shift has a discontinuity at shift day 12 since the bird did not sing at all that day. Also note how one bird in the −1 semitone shift group is at or slightly above zero by the end of the shift. This bird is the reason for the group not being statistically significantly below zero (this bird also had an extended baseline of 6 days; see *Materials and Methods*).

For 6-OHDA lesioned birds however, all birds decreased their pitch over time regardless of the direction of pitch shift through the headphones (Fig. 4b), similar to what we observed in lesioned birds with no pitch shifts (Fig. 3b). The +1 semitone shift group had a final pitch change of −0.38 ± 0.16 semitones (probability of resampled mean pitch greater than or equal to zero was p = 0.0040) while the −1 semitone shift group changed to a final pitch of −0.46 ± 0.19 semitones (probability of resampled mean pitch greater than or equal to zero was p = 0.0747) relative to the baseline (traces for individual birds are shown in Fig. 4-1b). The two groups were not statistically different from each other (probability of resampled mean pitch of +1 semitone shift group being greater than or equal to that of −1 semitone shift group was p = 0.26). We also compared each group to the no shift group and did not find statistically significant results (probability of resampled mean pitch of no shift group being greater than or equal to that of −1 semitone shift group was p = 0.62; probability of resampled mean pitch of no shift group being greater than or equal to that of +1 semitone shift group was p = 0.91). All statistical comparisons have been summarized in Table 1. Furthermore, when we quantified the adaptive change in pitch for this group, the final change in pitch was close to zero (gray trace, Fig. 4c). This suggests that following 6-OHDA lesions, birds do not respond adaptively to the auditory error. Instead, the birds seem to reduce their pitch over time regardless of the direction or presence of pitch-shifted auditory error. Note that as was mentioned above and shown in Table 1, there was not a statistically significant difference between the Lesioned −1 semitone shift group and zero. This was due to the fact that while birds subjected to the −1 semitone shift did reduce their pitch on average, a few syllables for each bird increased their pitch, resulting in a group effect that fell short of significance. Since our error quantification treats the contribution from each syllable equally, the effects of individual syllables add up resulting in a not statistically significant difference (see *Error Quantification* under *Materials and Methods*).

**Table 1:** Statistical tests summary. Results of statistical tests for ±1 semitone shift and 0 shift lesioned groups. The probabilities for each hypothesis are reported by testing the probability of the group on the left being greater than or equal to the various column headings. Blank spaces represent tests that either do not make sense to make or have been reported on another row.

| Hypothesis tested - Bayesian Probability of group on left being ≥ column heading (see *Hypothesis testing with Bootstrap* in *Materials and Methods*) | | | |
| --- | --- | --- | --- |
| Groups Compared | Zero | Lesioned +1 semitone shift | Lesioned -1 semitone shift |
| Lesioned 0 shift | **0.0029** | 0.91 | 0.62 |
| Lesioned +1 semitone shift | **0.0040** |  | 0.26 |
| Lesioned -1 semitone shift | 0.0747 |  |  |

Since the hierarchical bootstrapping as we have performed here to calculate statistical tests and standard errors has not been widely applied to such datasets in neuroscience previously, we also analyzed our data using hierarchical linear mixed models (LMMs) (Aarts et al., 2014; Aarts et al., 2015). LMMs have been widely applied to datasets involving large numbers of samples from a small number of subjects such as non-human primate studies (Arlet et al., 2015; Pleil et al., 2016) and rodent studies (Liang et al., 2015) or to analyze repeated measures or time series data (Wykes et al., 2012; Howe et al., 2013). Specifically, we built LMMs to test the effects of the shift condition while controlling bird identity and specific syllables within each bird as variable effects (see *Validating our Results with Linear Mixed Models* in *Materials and Methods*). For the unlesioned birds the linear mixed model revealed a strong effect of the shift condition (t = 7.17; p = 7.92 * 10^−13^) on final pitch at the end of the shift period. For the 6-OHDA lesioned birds, the effect of the shift condition (+1 semitone shift vs −1 semitone shift vs no shift) was not significant (t = 1.91; p = 0.056). Also, when we combined the shift groups and compared them to the no shift groups, the effect was not statistically significant (t = 1.47; p = 0.14). That these models give us the same statistically significant results as our bootstrapping procedure gives us an independent verification of our error calculation and statistics.

### 4. No correlations between lesion extent and changes in pitch

We measured the extent of 6OHDA lesions by quantifying the proportion of histological sections that fell below the 5^th^ percentile of section OD ratio for saline injected birds (see Methods). We can use this same threshold to obtain a rough metric of the lesion extent for each bird. Using this lesion extent, we computed correlations between the lesion extent and a variety of metrics of changes in pitch during the experiment (Table 1). However, we saw no significant correlations.

**Table 2:** Correlations between lesion extent and changes in song metrics. The lesion extent for each bird was defined as the proportion of sections with Optical Density (OD) ratio below the 5^th^ percentile of OD ratios for the population of saline injected birds. A Pearson’s correlation coefficient (sr) and the associated p-value is reported for this lesion extent versus changes in song metrics. Variances were computed across either three days of baseline or the final three days of the shift period.

| Lesion extent versus: | Pearson's correlation, r | Correlation significance, p |
| --- | --- | --- |
| Final pitch change | 0.4261 | 0.1466 |
| Baseline variance | 0.296 | 0.3261 |
| Final variance | -0.0498 | 0.8716 |
| Percent increase in variance | -0.4272 | 0.1454 |

### 5. Washout is impaired by dopamine depletion

Following the end of the shift period, we turned the pitch shift through the headphones back to zero and recorded the birds’ songs for an additional 6-7 days. During this period, birds without lesions typically revert their pitch back towards baseline levels (Sober and Brainard, 2009). Hence, we refer to this period as washout. We first collected washout data from the birds that had 6-OHDA lesions and headphones but no shifts. As stated earlier, by days 12 through 14 of the shift period, these birds had a mean pitch of −0.20 ± 0.13 semitones. By days 6 and 7 of the washout period, their pitch had changed to −0.34 ± 0.15 semitones (Fig. 5a; traces for individual birds are shown in Fig. 5-1a). The probability of the resampled mean pitch during the end of the shift period being greater than or equal to that during the end of the washout period was p = 0.67. Therefore, although the change was not statistically significant, the mean pitch did drop further during washout. In order to quantify how much the pitch changes in response to the end of the sensory perturbation (pitch shift), we subtracted the mean pitch for each syllable on the last day of pitch shift throughout the entire washout period and quantified the resulting deviation in pitch (Fig. 6a). This emphasizes the dynamics of how the pitch changes or Δ(Pitch) over time during washout in response to the end of the shift. The resulting change in pitch was found to be −0.12 ± 0.11 semitones (probability of resampled mean pitch greater than or equal to zero was p = 0.22).

**Figure 5:**
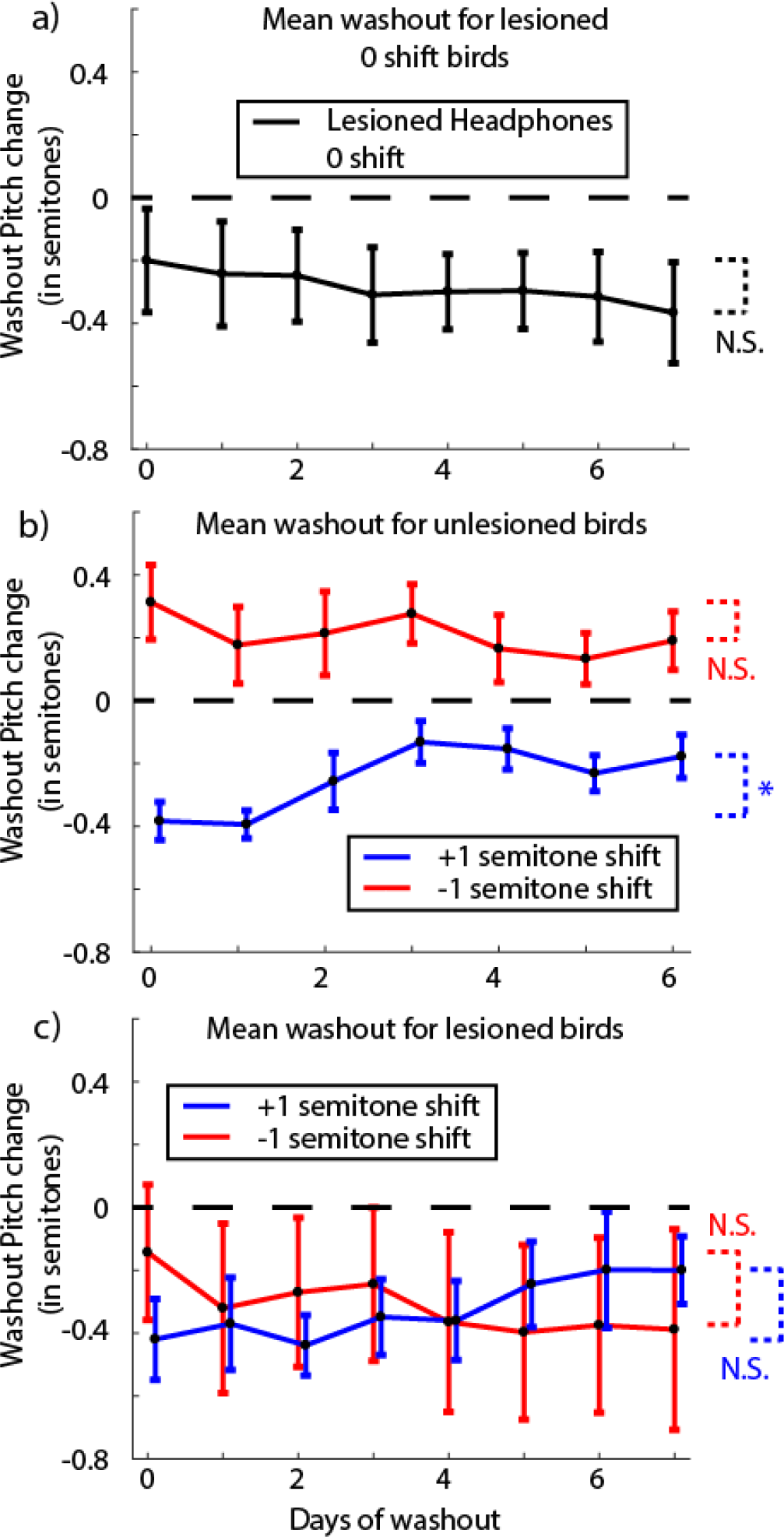
Analysis of change in pitch during washout for lesioned and unlesioned birds. a) Mean change in pitch during “washout” for lesioned birds with headphones but no pitch shift (N = 5 birds). Day 0 refers to the last day of the shift period. Pitch shift is turned off at the end of this day. Individual bird traces are shown in Figure 5-1a. b) Mean change in pitch during washout for unlesioned birds (N = 3 birds for each trace). Individual bird traces are shown in Figure 5-1b. c) Mean change in pitch during washout for 6-OHDA lesioned birds (N = 2 birds for each trace). The extremely large error bars are due in part to the bimodal nature of the data (see individual birds in Fig. 5-1c). The statistical tests check the last three days of the shift period against the last two days of washout with * representing a significant difference (p<0.05) and N.S. representing “not significant” (see Results for full tests).

**Figure 5-1:**
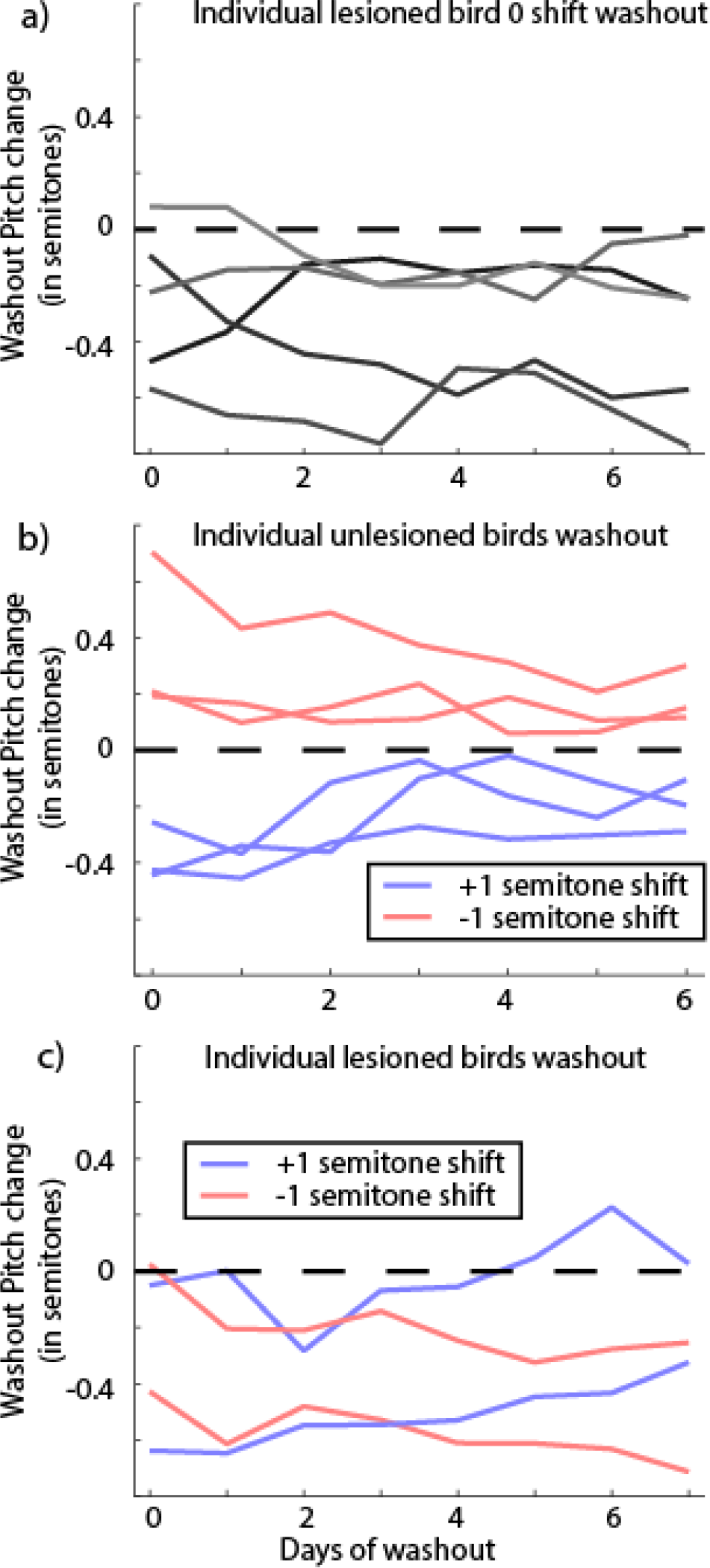
Washout traces for individual birds. a) Individual birds that had a 6-OHDA lesion, with headphones but no pitch shift. Each color is a separate bird. b) Washout traces for individual birds that were unlesioned and subjected to a ±1 semitone pitch shift. c) Washout traces for individual 6-OHDA lesioned birds subjected to a ±1 semitone pitch shift.

**Figure 6:**
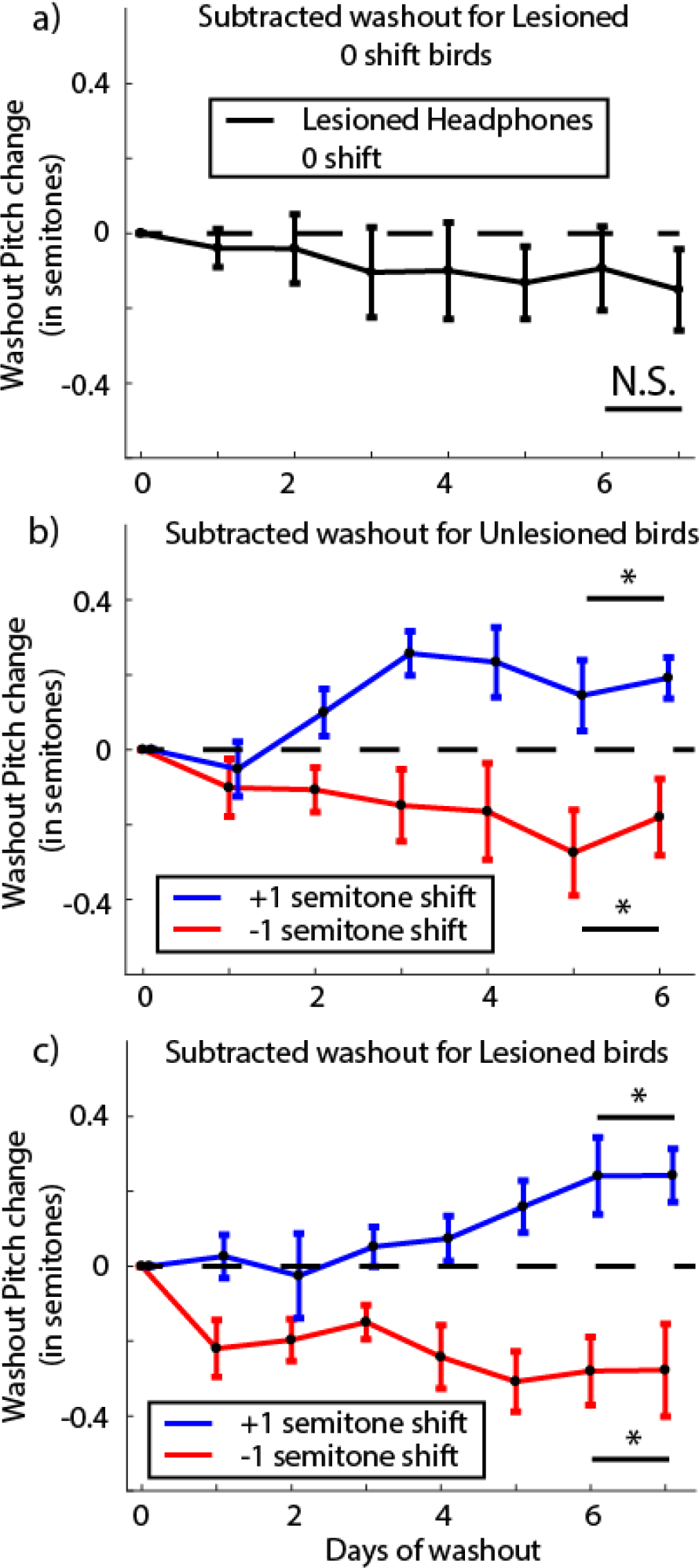
Results when measuring the dynamics of the change of pitch or Δ(Pitch) during washout by subtracting out the pitch change on the last day of shift through the washout period. Note that this figure shows the same data as Figure 5, but with pitch data plotted relative to the pitch on the final shift day rather than to the experiment’s baseline period as in Figure 5, a) Δ(Pitch) during washout for lesioned no shift birds (N = 5 birds). b) The same analysis as in a) for unlesioned birds subjected to ± 1 semitone shift (N = 3 birds each). c) The same analysis as in a) for lesioned birds subjected to ± 1 semitone shift (N = 2 birds each). The * and N.S. refer to a significant difference versus not respectively for each group compared to zero over the last two days of washout.

Unlesioned birds displayed a robust return to baseline following the end of the pitch shift period as shown in Figure 5b (see traces for individual birds in Fig. 5-1b). For birds subjected to a −1 semitone shift, they reduced their pitch from 0.36 ± 0.11 semitones at the end of shift to 0.17 ± 0.08 semitones during the last 2 days of washout (probability of mean resampled pitch during washout being greater than or equal to that at the end of shift was p = 0.08). Equivalently, birds subjected to a +1 semitone shift increased their pitch from −0.40 ± 0.07 semitones at the end of the shift period to −0.20 ± 0.05 semitones by the end of the washout period (probability of mean resampled pitch during washout being greater than or equal to that at the end of shift was p = 0.98). We also computed the dynamics underlying the Δ(Pitch) over time during the washout period by subtracting the pitch for each syllable on the last day of shift through the washout period (Fig. 6b). Birds subjected to a +1 semitone shift, having reduced their pitch during the shift increased their pitch during washout. The last 2 days of washout had a mean change relative to the last day of shift of 0.17 ± 0.07 semitones (probability of resampled mean pitch lesser than or equal to zero was p = 0.0003). Similarly, birds subjected to a −1 semitone shift reduced their pitch back towards baseline during washout by −0.22 ± 0.11 semitones relative to the last day of shift (probability of resampled mean pitch greater than or equal to zero was p = 0.0064).

For our 6-OHDA lesioned birds, only 4 out of 8 birds had data for 7 days of washout due to difficulties in keeping the headphones attached (2 each for upward and downward shifts). We repeated the analysis for washout for these birds as described above for lesioned no shift and unlesioned birds. First, the mean change in pitch from the last day of shift through the washout period is shown in Figure 5c. Birds subjected to a +1 semitone shift returned their pitch back towards baseline increasing their pitch from −0.31 ± 0.19 semitones at the end of the shift period to −0.20 ± 0.14 semitones by the end of the washout period (blue trace in Fig. 5c, probability of mean resampled pitch during washout being greater than or equal to that at the end of shift was p = 0.75). Contrary to expectations however, the birds subjected to a −1 semitone shift drifted further away from baseline reducing their pitch from −0.16 ± 0.22 semitones at the end of the shift to −0.38 ± 0.30 semitones by the end of the washout period (red trace in Fig. 5c, probability of mean resampled pitch during washout being greater than or equal to that at the end of shift was p = 0.35). The traces for individual birds are shown in Figure 5-1c.

Curiously, when we quantified the change in pitch in response to the end of the sensory perturbation subtracting the pitch change through the last day of shift through the washout period as before (i.e. measured the direction of pitch changes during washout, without considering the magnitude or direction of the pitch changes at the end of the shift period), the dynamics of the change in pitch was very similar to that seen in unlesioned birds (Fig. 6c). Lesioned birds subjected to a +1 semitone shift, averaging across the last 2 days of washout, shifted their pitch 0.24 ± 0.06 semitones with respect to the last day of shift (probability of resampled mean pitch lesser than or equal to zero was p = 0.0003). Lesioned birds subjected to a −1 semitone shift on the other hand, changed their pitch by −0.28 ± 0.11 semitones with respect to the last day of shift (probability of resampled mean pitch greater than or equal to zero was p = 0.0182). This result once again shows the dual effects we are observing following dopamine depletion. First, while not statistically significant, the pitch continued to drop for birds with unshifted auditory feedback. On the other hand, washout results between lesioned and unlesioned shift birds were very different in that washout is severely impaired in lesioned birds but confusingly followed the same dynamics for the Δ(Pitch) over time following the end of the pitch shifted auditory feedback.

## Discussion

Our results reveal two key effects of dopamine manipulation on the control of birdsong. First, all birds subjected to a 6-OHDA lesion of Area X displayed a drop in average vocal pitch which appeared between a week and two weeks post-lesion (Fig. 3b and Fig. 4b). Second, 6-OHDA lesioned birds displayed a severe deficit in sensorimotor learning as is evidenced by the lack of difference in response to a +1 or −1 semitone shift in pitch (Fig. 4b and gray trace in Fig. 4c).

While our primary finding seems to be one that implicates a role for dopamine in motor production, i.e., ability to produce higher pitched renditions of syllables in a bird’s repertoire, there is also a clear role for dopamine in learning the adaptive response to a sensory perturbation. It is true that when subjected to a +1 semitone pitch shift, there was no difference in mean change of pitch between lesioned (−0.38 ± 0.16 semitones) and unlesioned (−0.40 ± 0.07 semitones) birds (Fig. 4a and b, blue traces). However, when subjected to a −1 semitone pitch shift, while the adaptive response would be to raise their pitch, lesioned birds lowered their pitch (red trace in Fig. 4b). In addition, even for the lesioned birds subjected to a +1 semitone shift, their final change in pitch was not statistically different from the pitch drift seen in lesioned birds with no pitch shift (compare black trace in Fig. 3b with blue trace in Fig. 4b). This impairment in sensorimotor learning is reminiscent of deficits in learning in persons with Parkinson’s disease (Paquet et al., 2008; Mollaei et al., 2013) and rodent models of dopamine depletion in striatum and motor cortex (Shiotsuki et al., 2010; Hosp et al., 2011; Hosp and Luft, 2013). Hence our results suggest two factors at play, namely, motor production and sensorimotor learning. Disentangling these has been a hard problem in neuroscience (Beninger, 1983; Wise, 2004) since manipulations that affect motor learning also degrade motor production, complicating efforts to isolate learning mechanisms (Ungerstedt, 1968; Iancu et al., 2005; Cenci and Lundblad, 2007). Here, we isolated the lesions’ effects on motor production by including the lesioned no shift group.

We have previously reported that 6-OHDA lesions of Area X do not produce any changes in number of songs produced or in any general motor behavior (Hoffmann et al., 2016). We similarly did not observe any qualitative difference in song quality or motor behavior between lesioned birds reported in this study and the birds reported in the 2016 study except the systemic drop in average pitch of songs sung post-lesion. Note however that the lesioned birds reported in this study were recorded from for 2 to 3 weeks longer post-lesion than those from the 2016 study due to differences in time required to complete the behavioral experiments post-lesion. It therefore seems likely that this extended timeframe was necessary to observe the aforementioned pitch drop.

A reduction in motor “vigor” following dopamine depletion could explain the systemic drop in pitch we observed. Dopamine has been shown to be associated with vigor in humans and other mammalian systems (Niv et al., 2007; Beierholm et al., 2013; Panigrahi et al., 2015; Berke, 2018). Vigor has been characterized as motivation (Salamone et al., 2007; Salamone and Correa, 2012), speed of movements, or both (Mazzoni et al., 2007; Turner and Desmurget, 2010). In our experiments, we found that following 6-OHDA lesions of Area X the average pitch across all syllables for each bird dropped by roughly 11 to 13 days post-lesion. Higher pitched syllables require a combination of greater muscle activation and higher air sac pressure to be produced (Goller and Suthers, 1996; Elemans et al., 2008; Riede et al., 2010; Elemans et al., 2015) suggesting that higher pitched renditions of a particular syllable are more effortful to produce than lower pitched ones. We thus hypothesize that while unlesioned birds are capable of flexibly changing their pitch in a bidirectional fashion, dopamine lesioned birds will rarely increase their pitch due to the increased effort required to do so. A related observation supporting our interpretation of our results is that birds sing at an elevated pitch when singing directed songs to females (Sakata et al., 2008; Leblois et al., 2010). Since it has also been reported that dopamine levels in Area X are elevated during directed song (Sasaki et al., 2006), this fits with the overall trend in our results.

Studies that have targeted individual syllables for pitch changes following dopamine depletions have not reported a systemic drop in pitch post-lesion (Hoffmann et al., 2016; Hisey et al., 2018). Our study does not necessarily contradict these results since those studies reported a deficit in learning post-lesion by either combining upwards and downwards shifts (Hoffmann et al., 2016) or only driving pitch changes in one direction (Hisey et al., 2018). Additionally for the birds reported in this study, while the average pitch across all syllables for each bird dropped, some individual syllables did increase their pitch. Furthermore, as noted above the birds in the present study were recorded for a longer period of time post-lesion than those reported previously. The results from our washout data from the 6-OHDA lesioned birds are challenging to interpret. It is true that the lesioned birds subjected to a +1 semitone shift did return their pitch towards baseline and washout seemed to be unaffected for these birds (blue trace, Fig. 5c). Previous studies have reported that washout was not affected by dopamine depletion in tasks where birds shifted the pitch of a single syllable to avoid an aversive auditory cue (Hoffmann et al., 2016; Hisey et al., 2018). However, the birds subjected to a −1 semitone shift reduced their pitch resulting in their mean pitch moving further away from the baseline pitch (red trace, Fig. 5c). This suggests that washout is severely impaired in dopamine depleted birds. On the other hand, curiously, the change in pitch over time analyzed during washout in response to the end of the shift period was very similar between lesioned and unlesioned birds (compare Fig. 6b and 6c). We speculate that the lesion effects reported above could reflect either an inability to adaptively modulate motor output in response to error signals or from miscalculations in computing the error in the first place.

Adaptive sensorimotor learning in songbirds in response to induced auditory pitch shifts has been an effective paradigm to study the computational principles underlying sensorimotor learning (Sober and Brainard, 2009, 2012; Kelly and Sober, 2014). Bayesian inference works well to explain how unlesioned birds respond to auditory errors based on their prior experience of singing (Hahnloser and Narula, 2017; Zhou et al., 2018). However, since 6-OHDA lesioned birds exhibit drops in vocal pitch regardless of the direction of feedback pitch shift, any model that performs an adaptation to an error signal will fail to replicate the data without an additional mathematical mechanism to drive pitch downward in the presence of a reduced dopamine signal. One potential modification to the model would be to add a “relaxation state” into which the system relaxes in the absence of dopamine (Shadmehr and Arbib, 1992; Shadmehr and Mussa-Ivaldi, 1994). However, apart from the mean pitch, which did drop consistently across groups following 6-OHDA lesions, we did not find any other consistent relationships among other moments such as variance, skewness and kurtosis or overall probability distributions of produced pitch that could be used to constrain a revised Bayesian model to explain our results. Future work might therefore investigate the hypothesis that dopamine lesions disrupt sensorimotor learning by degrading the brain’s ability to perform Bayesian inference.

To conclude, our experiments show that dopamine plays a critical role in the brain’s ability to modulate vocal production in response to auditory errors. Future experiments will focus on disentangling specific roles for dopamine in sensorimotor learning by manipulating the dopamine signal at a faster temporal resolution. Results from such experiments could help fill gaps regarding the roles of tonic and phasic dopamine (Grace, 1991) for example and the timeline of error correction. Eventually, results from various such experiments can be used to impose mathematical constraints on a computational model detailing the quantitative role of dopamine in such sensorimotor learning.

## Acknowledgements

The work for this project was funded by the National Institutes of Health National Institute of Neurological Disorders and Stroke F31 NS100406, National Institutes of Health National Institute of Neurological Disorders and Stroke R01 NS084844, National Institutes of Health National Institute of Biomedical Imaging and Bioengineering R01EB022872, National Institutes of Health National Institute of Mental Health R01 MH115831-01, National Science Foundation 1456912, and Emory’s Udall Center of Excellence for Parkinson’s disease research. We would also like to thank David Hercules and Connor G Gallimore for their role in assisting Amanda L Jacob with tissue processing and imaging for some of the histology presented in this paper.

## References

Aarts E, Dolan CV, Verhage M, van der Sluis S (2015) Multilevel analysis quantifies variation in the experimental effect while optimizing power and preventing false positives. BMC Neurosci 16:94.

Aarts E, Verhage M, Veenvliet JV, Dolan CV, van der Sluis S (2014) A solution to dependency: using multilevel analysis to accommodate nested data. Nat Neurosci 17:491–496.

Arlet M, Jubin R, Masataka N, Lemasson AJBl (2015) Grooming-at-a-distance by exchanging calls in non-human primates. 11:20150711.

Balleine BW, O’doherty JP (2010) Human and rodent homologies in action control: corticostriatal determinants of goal-directed and habitual action. Neuropsychopharmacology 35:48.

Beierholm U, Guitart-Masip M, Economides M, Chowdhury R, Duzel E, Dolan R, Dayan P (2013) Dopamine modulates reward-related vigor. Neuropsychopharmacology 38:1495–1503.

Beninger RJ (1983) The role of dopamine in locomotor activity and learning. Brain Res 287:173–196.

Berke JD (2018) What does dopamine mean? Nat Neurosci 21:787–793.

Bottjer SWJJon (1993) The distribution of tyrosine hydroxylase immunoreactivity in the brains of male and female zebra finches. 24:51–69.

Brainard MS, Doupe AJ (2000) Interruption of a basal ganglia-forebrain circuit prevents plasticity of learned vocalizations. Nature 404:762–766.

Cenci MA, Lundblad M (2007) Ratings of L&-DOPA&-induced dyskinesia in the unilateral 6&-OHDA lesion model of Parkinson’s disease in rats and mice. Current protocols in Neuroscience 41:9.25. 21–29.25. 23.

Cooper JA, Sagar HJ, Jordan N, Harvey NS, Sullivan EV (1991) Cognitive impairment in early, untreated Parkinson’s disease and its relationship to motor disability. Brain 114 (Pt 5):2095–2122.

Crowley PH (1992) Resampling methods for computation-intensive data analysis in ecology and evolution. Annual Review of Ecology and Systematics 23:405–447.

Dubois B, Pillon BJJon (1996) Cognitive deficits in Parkinson’s disease. 244:2–8.

Efron B (1981) Nonparametric estimates of standard error: the jackknife, the bootstrap and other methods. Biometrika 68:589–599.

Efron B (1992) Bootstrap methods: another look at the jackknife. In: Breakthroughs in statistics, pp 569–593: Springer.

Efron B, Tibshirani RJ (1994) An introduction to the bootstrap: CRC press.

Elemans C, Rasmussen JH, Herbst CT, Düring DN, Zollinger SA, Brumm H, Srivastava K, Svane N, Ding M, Larsen ON (2015) Universal mechanisms of sound production and control in birds and mammals. Nature communications 6:8978.

Elemans CP, Mead AF, Rome LC, Goller F (2008) Superfast vocal muscles control song production in songbirds. PLoS One 3:e2581.

Gadagkar V, Puzerey PA, Chen R, Baird-Daniel E, Farhang AR, Goldberg JH (2016) Dopamine neurons encode performance error in singing birds. Science 354:1278–1282.

Galbraith S, Daniel JA, Vissel B (2010) A study of clustered data and approaches to its analysis. J Neurosci 30:10601–10608.

Glimcher PW (2011) Understanding dopamine and reinforcement learning: the dopamine reward prediction error hypothesis. Proc Natl Acad Sci U S A 108 Suppl 3:15647–15654.

Goller F, Suthers RA (1996) Role of syringeal muscles in controlling the phonology of bird song. J Neurophysiol 76:287–300.

Grace AA (1991) Phasic versus tonic dopamine release and the modulation of dopamine system responsivity: a hypothesis for the etiology of schizophrenia. Neuroscience 41:1–24.

Hahnloser RH, Narula G (2017) A Bayesian Account of Vocal Adaptation to Pitch-Shifted Auditory Feedback. PLoS One 12:e0169795.

Haith AM, Krakauer JW (2013) Model-based and model-free mechanisms of human motor learning. Adv Exp Med Biol 782:1–21.

Hisey E, Kearney MG, Mooney R (2018) A common neural circuit mechanism for internally guided and externally reinforced forms of motor learning. Nat Neurosci 21:589–597.

Hoffmann LA, Sober SJ (2014) Vocal generalization depends on gesture identity and sequence. J Neurosci 34:5564–5574.

Hoffmann LA, Kelly CW, Nicholson DA, Sober SJ (2012) A lightweight, headphones-based system for manipulating auditory feedback in songbirds. J Vis Exp:e50027.

Hoffmann LA, Saravanan V, Wood AN, He L, Sober SJ (2016) Dopaminergic Contributions to Vocal Learning. J Neurosci 36:2176–2189.

Hosp JA, Luft AR (2013) Dopaminergic meso-cortical projections to M1: role in motor learning and motor cortex plasticity. Frontiers in neurology 4:145.

Hosp JA, Pekanovic A, Rioult-Pedotti MS, Luft AR (2011) Dopaminergic projections from midbrain to primary motor cortex mediate motor skill learning. Journal of Neuroscience 31:2481–2487.

Howe WM, Berry AS, Francois J, Gilmour G, Carp JM, Tricklebank M, Lustig C, Sarter MJJoN (2013) Prefrontal cholinergic mechanisms instigating shifts from monitoring for cues to cue-guided performance: converging electrochemical and fMRI evidence from rats and humans. 33:8742–8752.

Iancu R, Mohapel P, Brundin P, Paul G (2005) Behavioral characterization of a unilateral 6-OHDA-lesion model of Parkinson’s disease in mice. Behavioural brain research 162:1–10.

Jankovic J (2008) Parkinson’s disease: clinical features and diagnosis. Journal of neurology, neurosurgery & psychiatry 79:368–376.

Jeon BS, Jackson-Lewis V, Burke RE (1995) 6-Hydroxydopamine lesion of the rat substantia nigra: time course and morphology of cell death. Neurodegeneration 4:131–137.

Kelly CW, Sober SJ (2014) A simple computational principle predicts vocal adaptation dynamics across age and error size. Front Integr Neurosci 8:75.

Kuebrich BD, Sober SJ (2015) Variations on a theme: Songbirds, variability, and sensorimotor error correction. Neuroscience 296:48–54.

Leblois A, Wendel BJ, Perkel DJ (2010) Striatal dopamine modulates basal ganglia output and regulates social context-dependent behavioral variability through D1 receptors. J Neurosci 30:5730–5743.

Lees AJ, Smith E (1983) Cognitive deficits in the early stages of Parkinson’s disease. Brain 106 (Pt 2):257–270.

Liang Z, Watson GD, Alloway KD, Lee G, Neuberger T, Zhang NJN (2015) Mapping the functional network of medial prefrontal cortex by combining optogenetics and fMRI in awake rats. 117:114–123.

Lipkind D, Marcus GF, Bemis DK, Sasahara K, Jacoby N, Takahasi M, Suzuki K, Feher O, Ravbar P, Okanoya K, Tchernichovski O (2013) Stepwise acquisition of vocal combinatorial capacity in songbirds and human infants. Nature 498:104–108.

Mandelblat-Cerf Y, Las L, Denisenko N, Fee MS (2014) A role for descending auditory cortical projections in songbird vocal learning. Elife 3.

Mazzoni P, Hristova A, Krakauer JW (2007) Why don’t we move faster? Parkinson’s disease, movement vigor, and implicit motivation. J Neurosci 27:7105–7116.

Mohan V, Morasso P, Metta GJN (2011) The distribution of rewards in sensorimotor maps acquired by cognitive robots through exploration. 74:3440–3455.

Mollaei F, Shiller DM, Gracco VL (2013) Sensorimotor adaptation of speech in Parkinson’s disease. Mov Disord 28:1668–1674.

Niv Y, Daw ND, Joel D, Dayan P (2007) Tonic dopamine: opportunity costs and the control of response vigor. Psychopharmacology (Berl) 191:507–520.

Panigrahi B, Martin KA, Li Y, Graves AR, Vollmer A, Olson L, Mensh BD, Karpova AY, Dudman JT (2015) Dopamine Is Required for the Neural Representation and Control of Movement Vigor. Cell 162:1418–1430.

Paquet F, Bedard MA, Levesque M, Tremblay PL, Lemay M, Blanchet PJ, Scherzer P, Chouinard S, Filion J (2008) Sensorimotor adaptation in Parkinson’s disease: evidence for a dopamine dependent remapping disturbance. Exp Brain Res 185:227–236.

Peh WY, Roberts TF, Mooney R (2015) Imaging auditory representations of song and syllables in populations of sensorimotor neurons essential to vocal communication. J Neurosci 35:5589–5605.

Person AL, Gale SD, Farries MA, Perkel DJ (2008) Organization of the songbird basal ganglia, including area X. J Comp Neurol 508:840–866.

Pleil KE, Helms CM, Sobus JR, Daunais JB, Grant KA, Kash TLJAb (2016) Effects of chronic alcohol consumption on neuronal function in the non&-human primate BNST. 21:1151–1167.

Redgrave P, Rodriguez M, Smith Y, Rodriguez-Oroz MC, Lehericy S, Bergman H, Agid Y, DeLong MR, Obeso JA (2010) Goal-directed and habitual control in the basal ganglia: implications for Parkinson’s disease. Nature Reviews Neuroscience 11:760.

Riede T, Fisher JH, Goller F (2010) Sexual dimorphism of the zebra finch syrinx indicates adaptation for high fundamental frequencies in males. PLoS One 5:e11368.

Sakata JT, Brainard MS (2006) Real-time contributions of auditory feedback to avian vocal motor control. J Neurosci 26:9619–9628.

Sakata JT, Brainard MS (2008) Online contributions of auditory feedback to neural activity in avian song control circuitry. J Neurosci 28:11378–11390.

Sakata JT, Hampton CM, Brainard MS (2008) Social modulation of sequence and syllable variability in adult birdsong. J Neurophysiol 99:1700–1711.

Salamone JD, Correa M (2012) The mysterious motivational functions of mesolimbic dopamine. Neuron 76:470–485.

Salamone JD, Correa M, Farrar A, Mingote SM (2007) Effort-related functions of nucleus accumbens dopamine and associated forebrain circuits. Psychopharmacology (Berl) 191:461–482.

Sasaki A, Sotnikova TD, Gainetdinov RR, Jarvis ED (2006) Social context-dependent singing-regulated dopamine. J Neurosci 26:9010–9014.

Scharff C, Nottebohm F (1991) A comparative study of the behavioral deficits following lesions of various parts of the zebra finch song system: implications for vocal learning. J Neurosci 11:2896–2913.

Schultz W, Dayan P, Montague PR (1997) A neural substrate of prediction and reward. Science 275:1593–1599.

Shadmehr R, Arbib MA (1992) A mathematical analysis of the force-stiffness characteristics of muscles in control of a single joint system. Biol Cybern 66:463–477.

Shadmehr R, Mussa-Ivaldi FA (1994) Adaptive representation of dynamics during learning of a motor task. J Neurosci 14:3208–3224.

Shiotsuki H, Yoshimi K, Shimo Y, Funayama M, Takamatsu Y, Ikeda K, Takahashi R, Kitazawa S, Hattori N (2010) A rotarod test for evaluation of motor skill learning. Journal of neuroscience methods 189:180–185.

Sober SJ, Brainard MS (2009) Adult birdsong is actively maintained by error correction. Nat Neurosci 12:927–931.

Sober SJ, Brainard MS (2012) Vocal learning is constrained by the statistics of sensorimotor experience. Proc Natl Acad Sci U S A 109:21099–21103.

Soha JA, Shimizu T, Doupe AJ (1996) Development of the catecholaminergic innervation of the song system of the male zebra finch. J Neurobiol 29:473–489.

Sohrabji F, Nordeen EJ, Nordeen KW (1990) Selective impairment of song learning following lesions of a forebrain nucleus in the juvenile zebra finch. Behav Neural Biol 53:51–63.

Tian LY, Brainard MS (2017) Discrete Circuits Support Generalized versus Context-Specific Vocal Learning in the Songbird. Neuron 96:1168–1177 e1165.

Turner RS, Desmurget M (2010) Basal ganglia contributions to motor control: a vigorous tutor. Curr Opin Neurobiol 20:704–716.

Ungerstedt U (1968) 6-Hydroxy-dopamine induced degeneration of central monoamine neurons. European journal of pharmacology 5:107–110.

Van Eycke YR, Allard J, Salmon I, Debeir O, Decaestecker C (2017) Image processing in digital pathology: an opportunity to solve inter-batch variability of immunohistochemical staining. Sci Rep 7:42964.

Wise RA (2004) Dopamine, learning and motivation. Nat Rev Neurosci 5:483–494.

Wolpert DM, Ghahramani Z, Jordan MI (1995) An internal model for sensorimotor integration. Science 269:1880–1882.

Wolpert DM, Diedrichsen J, Flanagan JR (2011) Principles of sensorimotor learning. Nat Rev Neurosci 12:739–751.

Wykes RC, Heeroma JH, Mantoan L, Zheng K, MacDonald DC, Deisseroth K, Hashemi KS, Walker MC, Schorge S, Kullmann DMJStm (2012) Optogenetic and potassium channel gene therapy in a rodent model of focal neocortical epilepsy.3004190.

Xiao L, Chattree G, Oscos FG, Cao M, Wanat MJ, Roberts TF (2018) A Basal Ganglia Circuit Sufficient to Guide Birdsong Learning. Neuron 98:208–221 e205.

Zhou B, Hofmann D, Pinkoviezky I, Sober SJ, Nemenman I (2018) Chance, long tails, and inference in a non-Gaussian, Bayesian theory of vocal learning in songbirds. Proc Natl Acad Sci U S A 115:E8538–E8546.

